# HRCHY-CytoCommunity identifies hierarchical tissue organization in cell-type spatial maps

**DOI:** 10.1101/2025.03.27.645631

**Authors:** Zekun Wang, Yafei Xu, Yuxuan Hu, Lin Gao

## Abstract

Diverse cell types within a tissue assemble into multicellular structures to shape the functions of the tissue. These structural modules typically comprise specialized subunits, each performing unique roles. We present HRCHY-CytoCommunity, a graph neural network-based framework that leverages differentiable graph pooling and graph pruning to identify hierarchical multicellular structures in cell phenotype-annotated single-cell spatial maps. HRCHY-CytoCommunity ensures the robustness of the result by employing a hierarchical majority voting strategy. To extend the utility of HRCHY-CytoCommunity in cross-sample comparative analysis, an additional cell-type enrichment-based clustering can be used to generate a unified set of nested multicellular structures across all tissue samples. Using single-cell-resolution spatial proteomics and transcriptomics data, we demonstrate that HRCHY-CytoCommunity outperforms the existing state-of-the-art method in discerning coarse-grained tissue compartments (TCs) and fine-grained tissue cellular neighborhoods (TCNs). To further validate the utility of multi-level nested multicellular structures, we apply HRCHY-CytoCommunity to breast cancer data with clinical information and reveal that large-scale TCs and small-scale TCNs enable the stratification of patients with different prognoses in a hierarchical manner.

## Introduction

The rapid advancement of spatial omics technologies has accelerated the in-depth study of tissue spatial structures. High-throughput sequencing techniques for spatial transcriptomics, such as Visium ^1^ and Slide-seq ^2, 3^, have the capability to detect the expression level of the entire transcriptome while preserving spatial information. Furthermore, imaging-based spatial transcriptomics techniques like MERFISH ^4^, and spatial proteomics techniques such as CODEX ^5^, IMC ^6^, and MIBI-TOF ^7^, can capture hundreds of RNAs or dozens of proteins within a single cell. In order to understand how tissue structures are related to their functions, researchers have introduced the concept of cellular neighborhoods or spatial domains, which serve as functional modules where various cell types interact and collaborate to maintain tissue functions ^5, 8, 9^. With an abundance of spatial omics data available, a lot of methods have been proposed for identifying cellular neighborhoods or spatial domains ^10-14^.

From a spatial perspective, a tissue typically consists of hierarchical multicellular structures, with large structures potentially containing small substructures of biological significance ^15, 16^. For instance, some tumor tissues can be coarsely divided into tumor compartments, immune compartments, and stroma compartments, which, in turn, can further be subdivided into substructures enriched with B cells and T cells, granulocytes, fibroblasts, and so on. As an example for illustrating the role of coarse-grained tissue structures, triple-negative breast tumors with separated tumor and immune compartments often result in good outcomes ^7^. As a representative fine-grained multicellular module, B cell and T cell-enriched tertiary lymphoid structure is known to serve as a favorable prognostic indicator for patients with various cancer types ^17^. In a broader sense, uncovering hierarchical multicellular structures holds significant importance for gaining deep insights into the assembly principles from individual cells to the entire tissue.

Currently, there are limited methods available for identifying hierarchical tissue structures. The state-of-the-art method, NeST ^18^, designed for spatial transcriptomics data, identifies nested hierarchical structures by finding coexpression hotspots, which are defined as groups of spatially colocalized cells that coexpress a set of genes. However, NeST uses gene expression features as inputs and thus may not be suitable for spatial omics data with single-cell resolution ^5, 6^, due to its limited number of available gene or protein expression features. Furthermore, the hierarchical structures identified by NeST may not cover all cells within the spatial omics data, and there may not be a clearly nested relationship between the identified different levels of structures. For instance, in some cases, cells belonging to the same fine-grained tissue structure may be assigned to different coarse-grained tissue structures, which may make it difficult to interpret the identities of these cells within the multi-level hierarchical structures. Additionally, NeST identifies spatially separated coexpression hotspots as distinct tissue structures, potentially impeding the discovery of structures with spatial discontinuous distribution.

We have recently established a tissue structure identification framework, CytoCommunity ^13^, which employs a graph neural network (GNN) model to learn a mapping directly from the cell type space to the tissue structure space, making it suitable for single-cell spatial omics data with limited gene or protein expression features and facilitating interpretation of functions of tissue structures. Building upon this general framework, we introduce HRCHY-CytoCommunity, a method dedicated for identifying hierarchical tissue structures based on cell types and their spatial locations. By leveraging differentiable graph pooling, graph pruning, and hierarchical majority voting, HRCHY-CytoCommunity enables simultaneous identification of robust tissue structures across various hierarchical levels that cover all cells within the spatial omics data and exhibit clearly nested relationship between them. This capability allows for a more nuanced understanding of how multicellular neighborhoods as building blocks form a tissue. For multi-sample analysis, HRCHY-CytoCommunity leverages cell-type enrichment-based clustering to align the nested tissue structures in different samples. For easy interpretation, we identified two-level hierarchical structures in all experiments in this study and defined the coarse- and fine-grained multicellular structures as the tissue compartment (TC) and the tissue cellular neighborhood (TCN), respectively. We benchmarked the performance of HRCHY-CytoCommunity on identifying TCs and TCNs using various single-cell spatial omics datasets. Using an additional breast cancer dataset with patient survival data, we also demonstrated the potential of identified hierarchical structures as prognostic indicators.

## Results

### Overview of HRCHY-CytoCommunity

HRCHY-CytoCommunity is an unsupervised learning framework that utilizes single-cell spatial maps, incorporating both cell type and spatial location information, to identify hierarchical multicellular structures in tissues. The framework consists of three modules: (1) a GNN-based module for soft hierarchical tissue structure assignment, (2) a consensus-driven module for hierarchical tissue structure ensemble, and (3) a clustering module that generates a unified hierarchical tissue structure set for cross-sample comparative analysis. Fig. 1 illustrates the identification process of hierarchical tissue structures at two levels. In the first module (Fig. 1a; Methods), HRCHY-CytoCommunity constructs a K-nearest-neighbor (KNN)-based cell-cell proximity graph, where nodes represent individual cells and edges reflect spatial proximity relationship among cells. Each node is equipped with an attribute vector encoding cell type information. Through a graph convolution layer and a fully connected layer, the framework transforms these cell-type attribute vectors into soft fine-grained tissue structure assignment vectors, where each dimension represents the probability of belonging to a specific fine-grained tissue structure. Then, HRCHY-CytoCommunity applies a graph pooling layer to generate a coarsened completed graph, where pooled nodes correspond to fine-grained tissue structures and edge weights quantify inter-structural connectivity strength. To mitigate potential over-smoothing effect from message passing during graph convolution, HRCHY-CytoCommunity implements edge pruning on the coarsened completed graph. Subsequent employment of graph convolution and fully connected layers on this pruned graph yields soft coarse-grained tissue structure assignments for the pooled nodes. The learning process is guided by two graph minimum cut (MinCut)-based loss functions ^19^, separately optimizing fine-grained and coarse-grained tissue structure assignments. These loss functions are combined through linear combination to form the overall optimization objective.

**Fig. 1.**
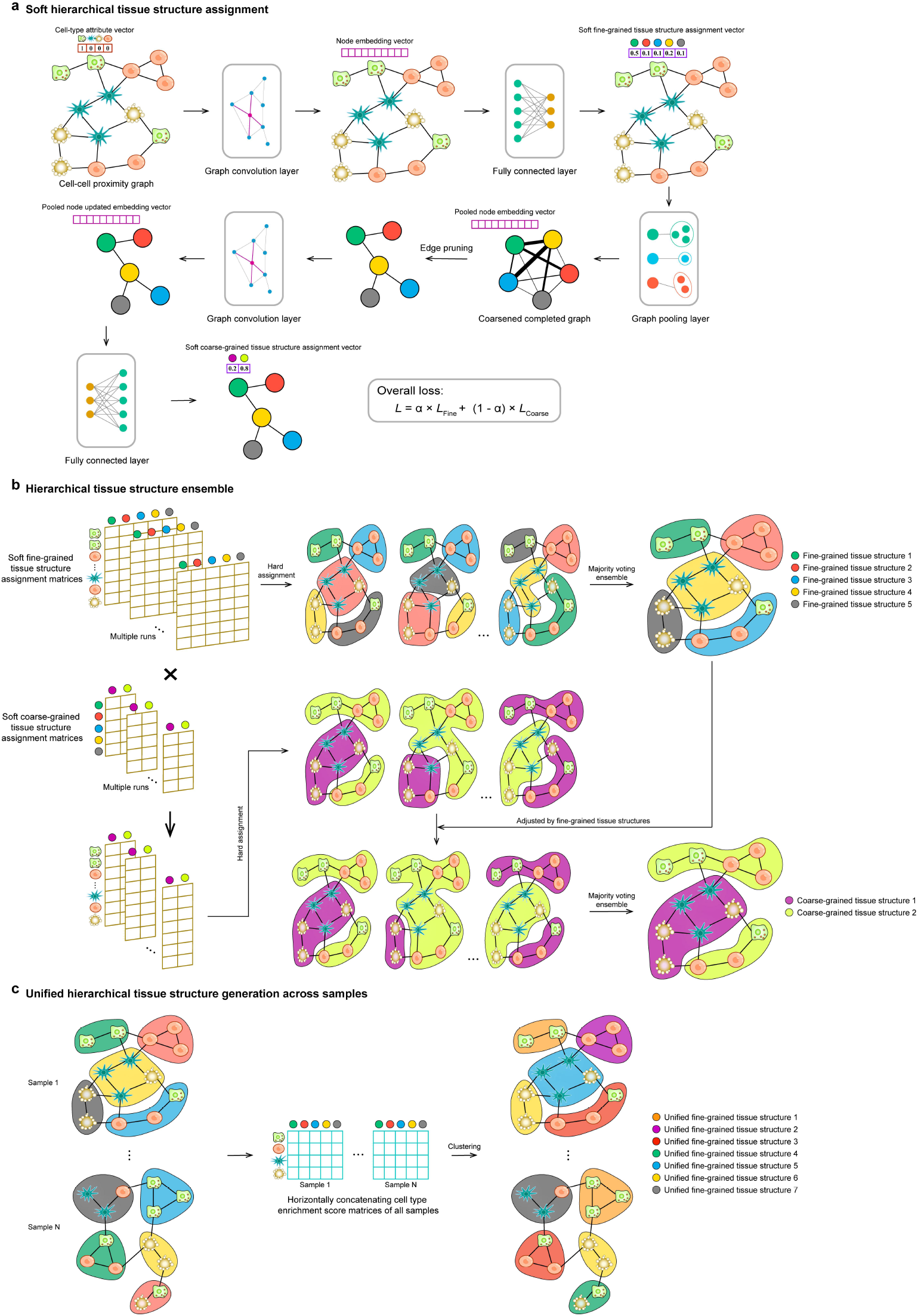
Overview of the HRCHY-CytoCommunity framework. The hierarchical tissue structure identification is modeled as a hierarchical community detection problem on a cell-cell proximity graph with node attributes. The HRCHY-CytoCommunity framework includes three modules: **(a)** a soft hierarchical tissue structure assignment module, **(b)** a hierarchical tissue structure ensemble module and **(c)** a unified tissue structure generation module used for cross-sample comparative analysis. **(a)** In the first module, a cell-cell proximity graph is constructed based on spatial relationships among cells, with nodes representing cells and one-hot encoded attributes indicating cell types. A graph convolution layer and a fully connected layer of the GNN transform cell type attributes into soft fine-grained tissue structure assignments. Subsequently, a graph pooling layer is used to generate a coarsened graph where each pooled node represents a fine-grained tissue structure. Edge pruning is used to mitigate over-smoothing effect from message passing. Another graph convolution and fully connected layers are utilized to generate the soft coarse-grained tissue structure assignment vector of each node, representing the probability that it is assigned to each coarse-grained tissue structure. The graph minimum cut (MinCut)-based loss function is used to learn fine-grained and coarse-grained tissue structure assignments. The overall loss function is a linear combination of two MinCut-based loss functions, employed for learning hierarchical tissue structure assignment. **(b)** In the hierarchical tissue structure ensemble module, a hierarchical majority-voting strategy is used to alleviate the instability issue of graph partitioning based on GNN and ensure the clearly nested relationship between coarse-grained and fine-grained tissue structures. First, the soft hierarchical tissue structure assignment module is run several times to obtain multiple soft fine-grained and coarse-grained tissue structure assignment matrices. Subsequently, we multiply the two sets of matrices to obtain a set of matrices representing the probabilities of assigning cells (rows) to coarse-grained tissue structures (columns). Next, for each matrix representing the probabilities that cells assigned to tissue structures, the hard assignment is conducted by assigning the cell (row) to the tissue structure (column) with the highest probability. Subsequently, the majority-voting strategy is used for all hard fine-grained tissue structure assignments to obtain robust fine-grained tissue structures. Each hard coarse-grained tissue structure assignment is adjusted by making it incorporate one or several complete fine-grained tissue structures. Finally, the majority-voting strategy is used for all adjusted hard coarse-grained tissue structure assignments to obtain robust coarse-grained tissue structures. **(c)** A clustering module that generates a unified hierarchical tissue structure set for cross-sample comparative analysis. The cell type enrichment score matrices of all samples are horizontally concatenated, and a clustering algorithm is used to generate a unified hierarchical tissue structure set for cross-sample comparative analysis.

The second module (Fig. 1b; Methods) implements an ensemble strategy to enhance the robustness of hierarchical tissue structure identification. The soft hierarchical tissue structure assignment module is executed multiple times, followed by a hierarchical majority-voting procedure that integrates these soft tissue structure assignments into a consensus solution. This ensemble approach effectively addresses the inherent instability of GNN-based graph partitioning while maintaining a clearly nested relationship between coarse-grained and fine-grained tissue structures.

Since HRCHY-CytoCommunity operates as an unsupervised learning framework optimized for single-sample analysis, cross-sample tissue structure comparison presents a significant computational challenge. To overcome this limitation, HRCHY-CytoCommunity incorporates a third module specifically designed for cross-sample tissue structure alignment (Fig.1c; Methods). This module first calculates cell type enrichment scores for (coarse- or fine-grained) tissue structures across all samples and concatenates the cell type enrichment score matrices into a unified feature matrix. Then, a clustering algorithm is applied to this composite feature matrix to cluster tissue structures from different samples into a unified (coarse- or fine-grained) tissue structure set. Notably, the resulting number of unified hierarchical tissue structures may exceed the pre-defined structure count, as this approach preserves sample-specific multicellular structures while identifying shared architectural patterns.

### Hierarchical tissue structures in the mouse spleen

To assess the ability of HRCHY-CytoCommunity to identify large-scale coarse-grained tissue structures, we applied it to a spatial proteomics dataset of mouse spleen generated obtained through the Co-Detection by Indexing (CODEX) technology ^20^. For benchmarking, we compared it with NeST ^18^, the existing state-of-the-art method for identifying nested hierarchical structures. This dataset comprises three healthy mouse spleen samples, named as “BALBc-1”, “BALBc-2”, and “BALBc-3”. These samples were annotated into 27 distinct cell types (Fig.2a, first column) and two major spleen compartments: the red pulp and the white pulp, serving as the gold standard for coarse-grained TCs (Fig.2a, second column).

**Fig. 2.**
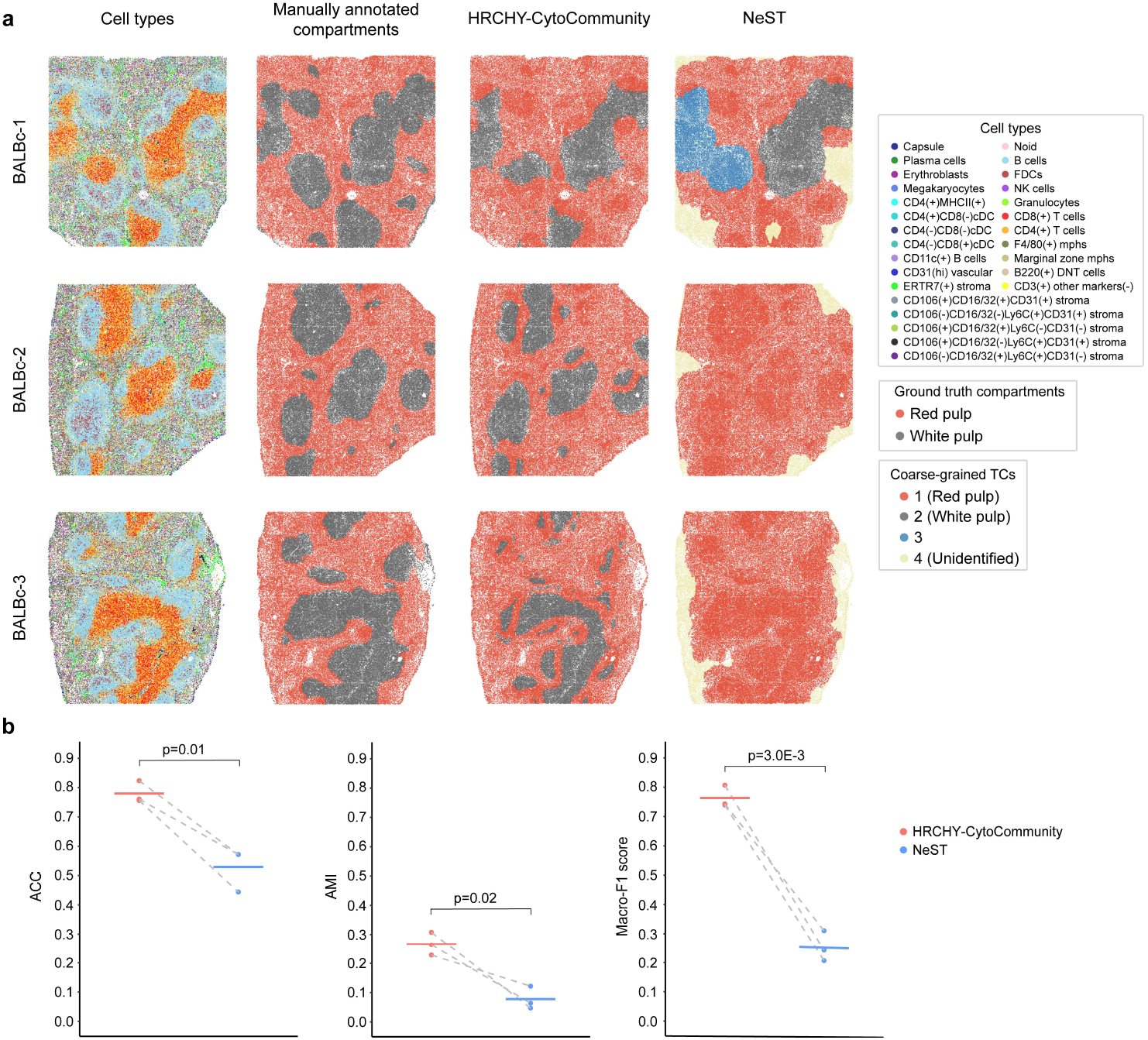
Evaluation of HRCHY-CytoCommunity’s performance on the mouse spleen CODEX dataset. **(a)** Single-cell spatial images of healthy mouse spleen samples obtained using CODEX technology. The cells are colored according to cell types (first column), manually annotated compartments (second column), and the coarse-grained TCs identified by HRCHY-CytoCommunity (third column) and NeST (fourth column). Unidentified indicates that the cells were not assigned to any TCs. **(b)** Accuracy (ACC), Adjusted Mutual Information (AMI) and Macro-F1 scores calculated using manually annotated compartments. Each point corresponds to the performance on an individual image, with horizontal bars indicating the mean performance across all images. Points from the same images are connected by grey dashed lines. *P*-values were calculated using a one-sided paired t-test.

We pre-defined the number of coarse-grained TC structures to two and the number of fine-grained TCN structures to seven, identifying the hierarchical tissue structures of three mouse spleen samples. We found that for all samples, HRCHY-CytoCommunity identified two TCs, corresponding to the red pulp and the white pulp (Fig.2a, third column). Notably, despite the white pulp was a spatially discontinuous structure, HRCHY-CytoCommunity was able to accurately identify it. In contrast, NeST identified nested hierarchical structures starting from single-gene hotspots ^18^, thus some cells were not assigned to any TCs (Fig.2a, fourth column, unidentified). For the BALBc-1 sample, NeST identified the spatially separated white pulp as two distinct TCs. For the BALBc-2 and BALBc-3 samples, NeST only identified the red pulp, possibly because the limited features of the dataset prevented the identification of the white pulp (Fig.2a, fourth column). To quantify the performance of identifying TCs, we measured the concordance between the predicted TCs and the manually annotated compartments using three evaluation metrics: Accuracy (ACC), adjusted mutual information (AMI), and Macro-F1 score (Methods). The results demonstrated that HRCHY-CytoCommunity achieved superior performance across all three samples (paired t-test p-values < 0.05; Fig.2b).

Furthermore, the coarse-grained red pulp compartment could be further divided into two fine-grained substructures, red pulp and marginal zone; while the coarse-grained white pulp compartment could be divided into B-cell zone and periarteriolar lymphoid sheath (PALS) ^20^ (Supplementary Fig.1, second column). For the TCNs identified by HRCHY-CytoCommunity, we found that TCN-1, TCN-2 and TCN-8 corresponded to the red pulp; TCN-5 and TCN-6 corresponded to the marginal region; TCN-4 corresponded to the B-zone; TCN-3 and TCN-7 corresponded to the PALS (Supplementary Fig.1, third column). In contrast, the TCNs identified by NeST had a poor correspondence with the manually annotated compartments (Supplementary Fig.1, fourth column). In conclusion, compared with the existing state-of-the-art hierarchical tissue structure identification method, HRCHY-CytoCommunity is better able to identify coarse-grained red pulp and white pulp compartments, while the identified fine-grained TCNs can better correspond to the known biological structures.

### Hierarchical tissue structures in the mouse hypothalamic preoptic region

To further assess the capability of HRCHY-CytoCommunity in identifying smaller fine-grained TCNs, we applied it to a spatial transcriptomics dataset of the healthy mouse hypothalamic preoptic region. This dataset was generated using the Multiplexed Error-Robust Fluorescence in situ Hybridization (MERFISH) technology ^4^, which measures the expression of 155 genes. It includes samples from five brain regions, named “Bregma-0.04”, “Bregma-0.14”, “Bregma+0.06”, “Bregma+0.16”, and “Bregma+0.26” based on their relative distances to the bregma ^4^ (Fig.3a, first column). The dataset encompasses 17 hypothalamic nuclei regions, which were delineated in the original study through manual inspection of well-characterized brain histology ^4, 21^ (Fig.3a, second column). We previously established a reference for fine-grained TCNs by manually mapping these nuclei onto single-cell spatial maps and annotating all cells accordingly (Fig.3a, third column).

**Fig. 3.**
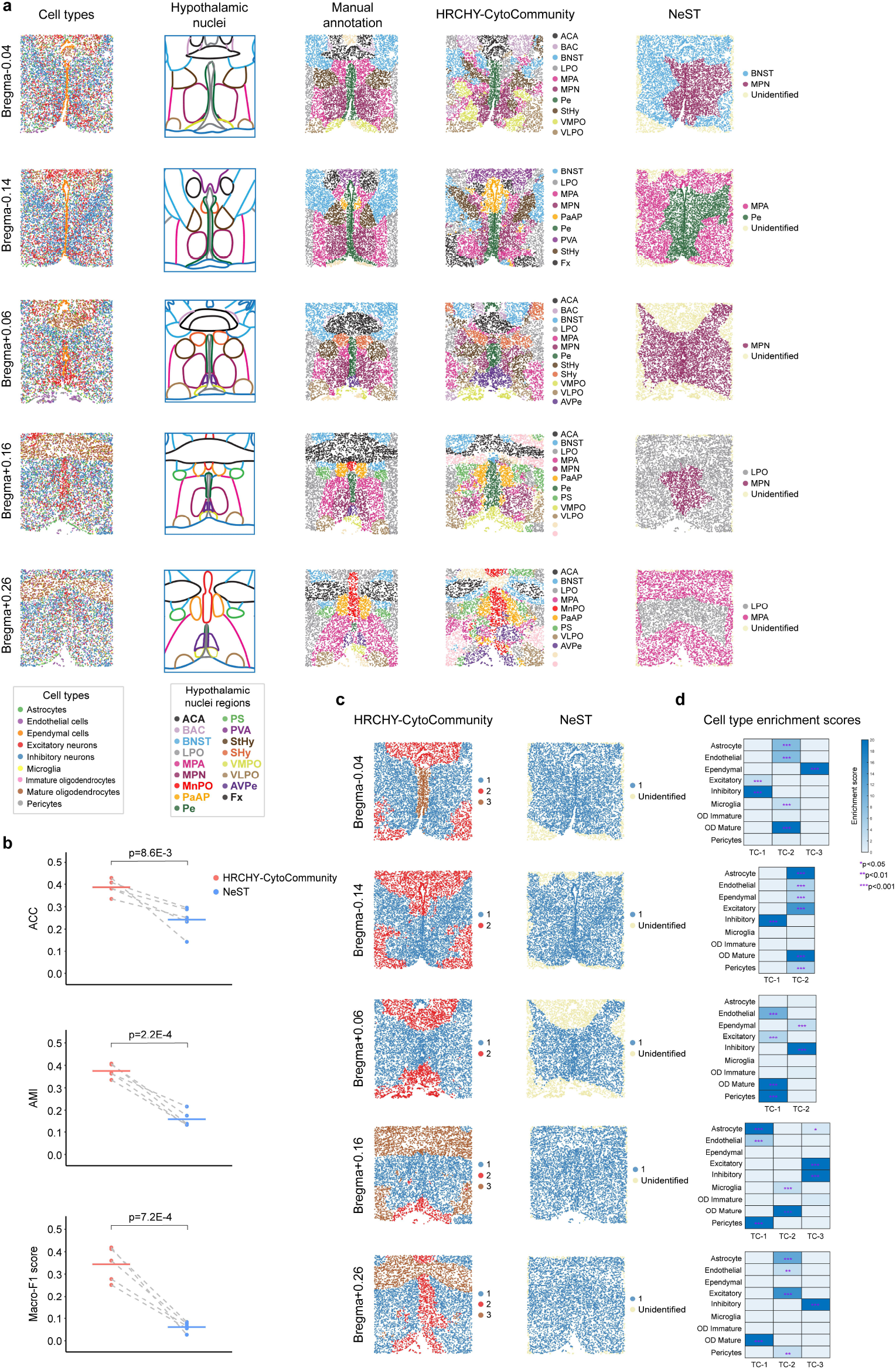
Evaluation of HRCHY-CytoCommunity’s performance on the mouse hypothalamic preoptic region MERFISH dataset. **(a)** Single-cell spatial images of mouse hypothalamic preoptic region obtained using MERFISH technology. The cells are colored according to cell types (first column), hypothalamic nuclei (second column), manual annotation (third column) and the fine-grained TCNs identified by HRCHY-CytoCommunity (fourth column) and NeST (fifth column). Unidentified indicates that the cells were not assigned to any TCNs. **(b)** Accuracy (ACC), Adjusted Mutual Information (AMI) and Macro-F1 scores calculated using manually annotation. Each point corresponds to the performance on an individual image, with horizontal bars indicating the mean performance across all images. Points from the same images are connected by grey dashed lines. *P*-values were calculated using a one-sided paired t-test. **(c)** Single-cell spatial images where cells are colored based on the coarse-grained TCs identified by HRCHY-CytoCommunity (first column) and NeST (second column). Unidentified indicates that the cells were not assigned to any TCs. **(d)** Heatmaps showing the average enrichment scores for each cell type within each coarse-grained TC across all images. The enrichment score for each cell type is calculated by −log_10_(*P*-value). *P*-values were derived using the hypergeometric test and subsequently adjusted with Benjamini-Hochberg procedure.

We pre-defined the number of fine-grained TCN structures to match the number of hypothalamic nuclei regions and the number of coarse-grained TC structures to three. We found that HRCHY-CytoCommunity successfully identified multiple symmetric TCNs consistent with manually delineated nuclei (Fig.3a, fourth column). For instance, BNST (Bed Nucleus of the Stria Terminalis), LPO (Lateral Preoptic Area) and MPA (Medial Preoptic Area) were consistently identified across all five samples. Larger regions, such as ACA (Anterior Commissure, Anterior Part) and MPN (Medial Preoptic Nucleus), were identified in several samples, while smaller regions, including PS (Parastrial Nucleus) and AVPe (Anteroventral Periventricular Nucleus), were also found in some samples. Additionally, unique structures like PVA (Paraventricular Nucleus, Anterior Part) and Fx (Fornix) were identified only in the “Bregma-0.14” sample, while SHy (Suprachiasmatic nucleus shell) was unique to the “Bregma+0.06” sample. In contrast, NeST identified only one or two TCNs in each sample (Fig.3a, fifth column), which differed from the manually delineated nuclei.

NeST performed poorer than HRCHY-CytoCommunity probably because it used gene expression features as inputs, and it did not specify the number of tissue structures in advance. Moreover, NeST was unable to identify spatially symmetric and discontinuously distributed parts of the same nuclear structure as the same tissue structure. Quantitatively, HRCHY-CytoCommunity demonstrated significantly higher ACC scores, AMI scores and Macro-F1 scores compared to NeST (paired t-test p-values < 0.01; Fig.3b).

Subsequently, we analyzed the coarse-grained TC structures of these five samples identified by HRCHY-CytoCommunity. For each sample, HRCHY-CytoCommunity identified 2-3 TCs, while NeST only identified one TC (Fig.3c). Notably, for the samples “Bregma-0.14” and “Bregma+0.06”, although the number of TCs was pre-defined to three, HRCHY-CytoCommunity leveraged its deep learning module to automatically determine two TCs based on the features of the samples (Methods). We calculated the cell type enrichment scores (Methods) of the TCs identified by HRCHY-CytoCommunity in each sample and found that different TCs were enriched for different types of cells (Fig.3d). In the “Bregma-0.04” and “Bregma+0.16” samples, we respectively identified a TC enriched with neuron cells and two TCs enriched with non-neuronal cells. While in the “Bregma-0.14”, “Bregma+0.06” and “Bregma+0.26” samples, we respectively identified a TC enriched with inhibitory neuron cells and one or two TCs enriched with excitatory neuron cells and non-neuronal cells. From this analysis it was observed that neuronal cells and non-neuronal cells can respectively form structures with different functions, and non-neuronal cells interact differently with excitatory neuron cells and inhibitory neuron cells, which is consistent with the prior knowledge that non-neuronal cells play a role in modulating neuronal excitability in the physiological and pathophysiological processes of the brain.^22^.

### Comparative analysis of hierarchical tissue structures in triple-negative breast cancer

To further evaluate the performance of HRCHY-CytoCommunity, we applied it to a spatial proteomics dataset comprising 41 triple-negative breast cancer (TNBC) patients, obtained using the Multiplexed Ion Beam Imaging by Time-Of-Flight (MIBI-TOF) technology ^7^. Within this dataset, we selected 15 compartmentalized tumor samples, where immune cells were spatially separated from neoplastic cells ^7^, and compared with NeST.

We pre-defined the number of coarse-grained TC structures to two and the number of fine-grained TCN structures to 15. HRCHY-CytoCommunity successfully identified two TCs in most samples with one dominated by immune cells and the other one dominated by neoplastic cells. Such result is consistent with spatial distribution patterns of cell types for patients with compartmentalized tumors. (Fig.4a; Supplementary Fig.2). In contrast, for patient 4 and 9, NeST only identified one major TC, possibly because the limited features of the dataset prevented simultaneous identification of both tumor and immune TCs (Fig.4a). For patient 28 where the tumor cell population and immune cell population are spatially distributed contiguously, both HRCHY-CytoCommunity and NeST were able to effectively identify TCs (Fig.4a, second row). Conversely, for patient 32 where the tumor cell population and immune cell population are spatially distributed discretely, HRCHY-CytoCommunity successfully identified them as separate tumor and immune TCs, while NeST identified them as five TCs (Fig.4a, fourth row).

**Fig. 4.**
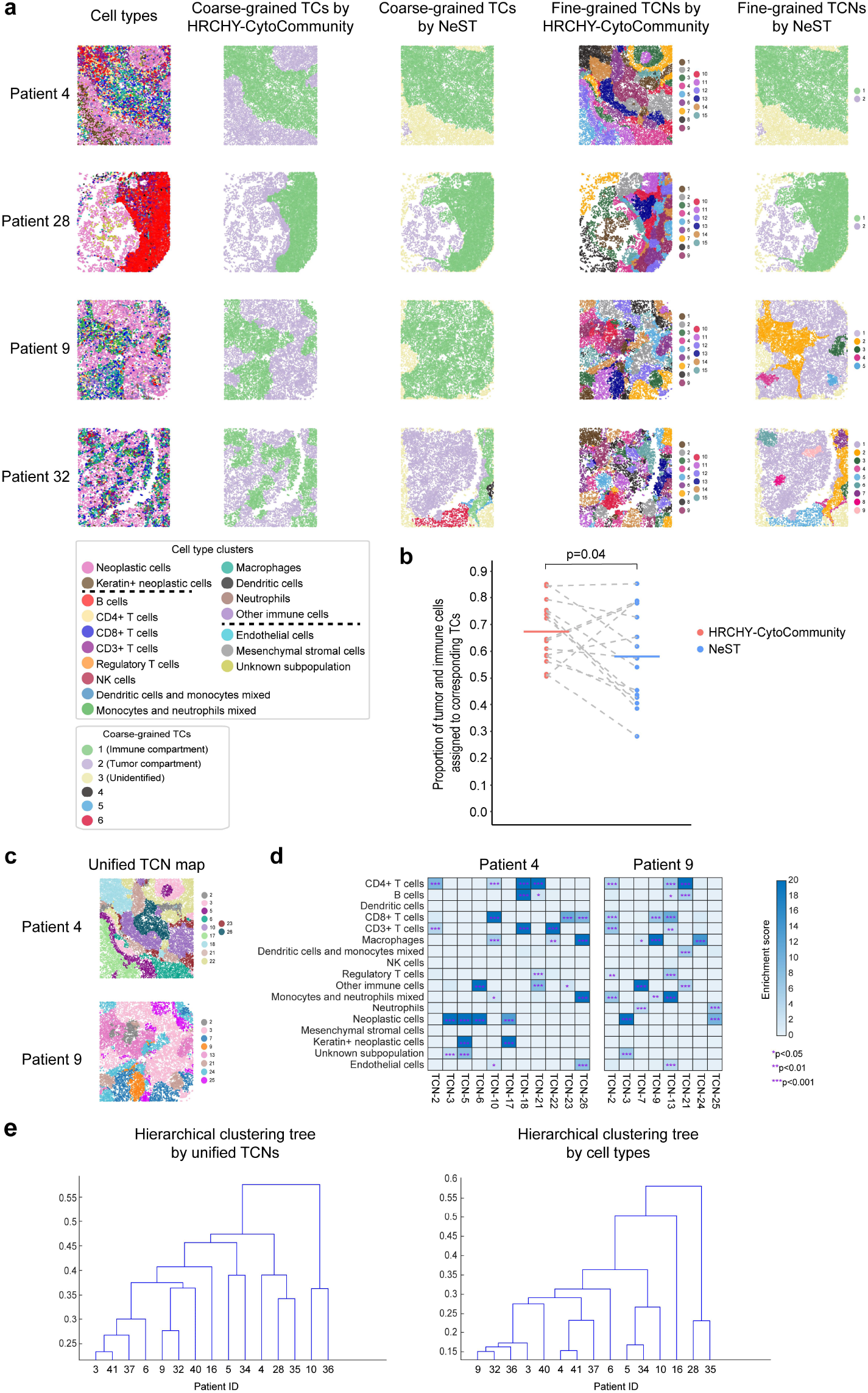
Evaluation of HRCHY-CytoCommunity’s performance on the TNBC MIBI-TOF dataset. **(a)** Representative single-cell images of the compartmentalized tumors from TNBC patients obtained using MIBI-TOF technology. The cells are colored according to cell types (first column), the coarse-grained TCs identified by HRCHY-CytoCommunity (second column) and NeST (third column), and the fine-grained TCNs identified by HRCHY-CytoCommunity (fourth column) and NeST (fifth column). Unidentified indicates that the cells were not assigned to any TCs. **(b)** Proportion of tumor and immune cells assigned to corresponding TCs. Each point corresponds to the performance on an individual image, with horizontal bars indicating the mean performance across all images. Points from the same images are connected by grey dashed lines. *P*-value was calculated using a one-sided paired t-test. **(c)** Unified TCN maps of patient 4 and patient 9. The cells are colored according to the unified TCNs. **(d)** Heatmaps showing the average enrichment scores for each cell type within each unified TCN in patient 4 and patient 9. The enrichment score for each cell type is calculated by −log_10_(*P*-value). *P*-values were derived using the hypergeometric test and subsequently adjusted with Benjamini-Hochberg procedure. **(e)** Hierarchical clustering trees by unified TCNs (left) and cell types (right).

To quantify the performance of TC identification, we calculated the proportion of neoplastic and immune cells that were correctly assigned to neoplastic and immune cell-dominated TCs. For those cells belonging to both tumor and immune TCs in the NeST results, we assigned them to the TC containing a greater proportion of cells (Methods). We found that HRCHY-CytoCommunity performs better across the 15 patients (paired t-test p-value = 0.04; Fig.2b). However, for patient samples with large scale differences between immune and tumor compartments, such as patient 35, 37, and 41, NeST performed better due to not being influenced by the scale of structures, while HRCHY-CytoCommunity performed worse due to minimizing the orthogonality loss term (Supplementary Fig.2; Methods). Nevertheless, with prior knowledge of this situation, setting an appropriate threshold for the orthogonality loss term could significantly improve the results (paired t-test p-value = 0.04; Supplementary Fig.3; Methods).

While compartmentalized tumors can be broadly divided into tumor TCs and immune TCs (Fig. 4a, second column), the fine-grained TCN analysis can reveal deeper heterogeneity within these tumors. To enable cross-sample comparison, we combined cell type enrichment score matrices across patient cohorts (Methods) and performed TCN clustering. This identified 26 unified TCNs that captured both conserved and patient-specific biological features (Fig.4c; Supplementary Fig.4). For example, the tumor TCs of both patient 4 and patient 9 contain the unified TCN-3 enriched with neoplastic cells and the immune TCs of these two patients contain the unified TCN-21 enriched with CD4+ T cells and B cells. However, CD3+ T cell and macrophage-enriched unified TCN-22 was only found in the immune TC of patient 4, but not in patient 9. In contrast, the unified TCN-25 enriched with neoplastic cells mixed with neutrophils was only found in the tumor TC of patient 9, but not in patient 4 (Fig.4d). These findings demonstrate that spatially organized tumors maintain both universal and distinct multicellular structures across individuals.

We further stratified patients with compartmentalized tumors using hierarchical clustering of unified TCN profiles. While some patients exhibited similar unified TCNs (such as patient 9 and patient 32), others showed noticed differences in the composition of unified TCNs (such as patient 9 and patient 10) (Fig.4e, left; Fig.4c; Supplementary Fig.4). To demonstrate that this heterogeneity does not solely arise from differences in cell types, hierarchical clustering was also conducted using cell types as features. The results revealed that although some patients (such as patient 4 and patient 41) had similar compositions of cell types, there were differences in the composition of their unified TCNs (Fig.4e; Fig.4c; Supplementary Fig.4). Just as the analysis of coarse-grained TCs can distinguish patients into compartmentalized tumor patients and mixed tumor patients ^7^, the analysis of fine-grained TCNs offers potential for further subclassifying these patients.

### Hierarchical tissue structure-based stratification of breast cancer patients

To validate the biological significance and clinical relevance of the hierarchical tissue structures identified by HRCHY-CytoCommunity, we applied these structures to stratify patients in a breast cancer spatial proteomics dataset generated using imaging mass cytometry (IMC) technology ^6^. The dataset consists of 252 breast cancer patients, with 179 surviving and 73 deceased. We pre-defined the number of coarse-grained TC structures to two and the number of fine-grained TCN structures to seven to identify the hierarchical tissue structures of each patient sample. Taking the patient with “PID = 11” as an example, each TC fully encompassed one or multiple TCNs, aligning with the spatial distribution patterns of cell types (Fig.5a).

**Fig. 5.**
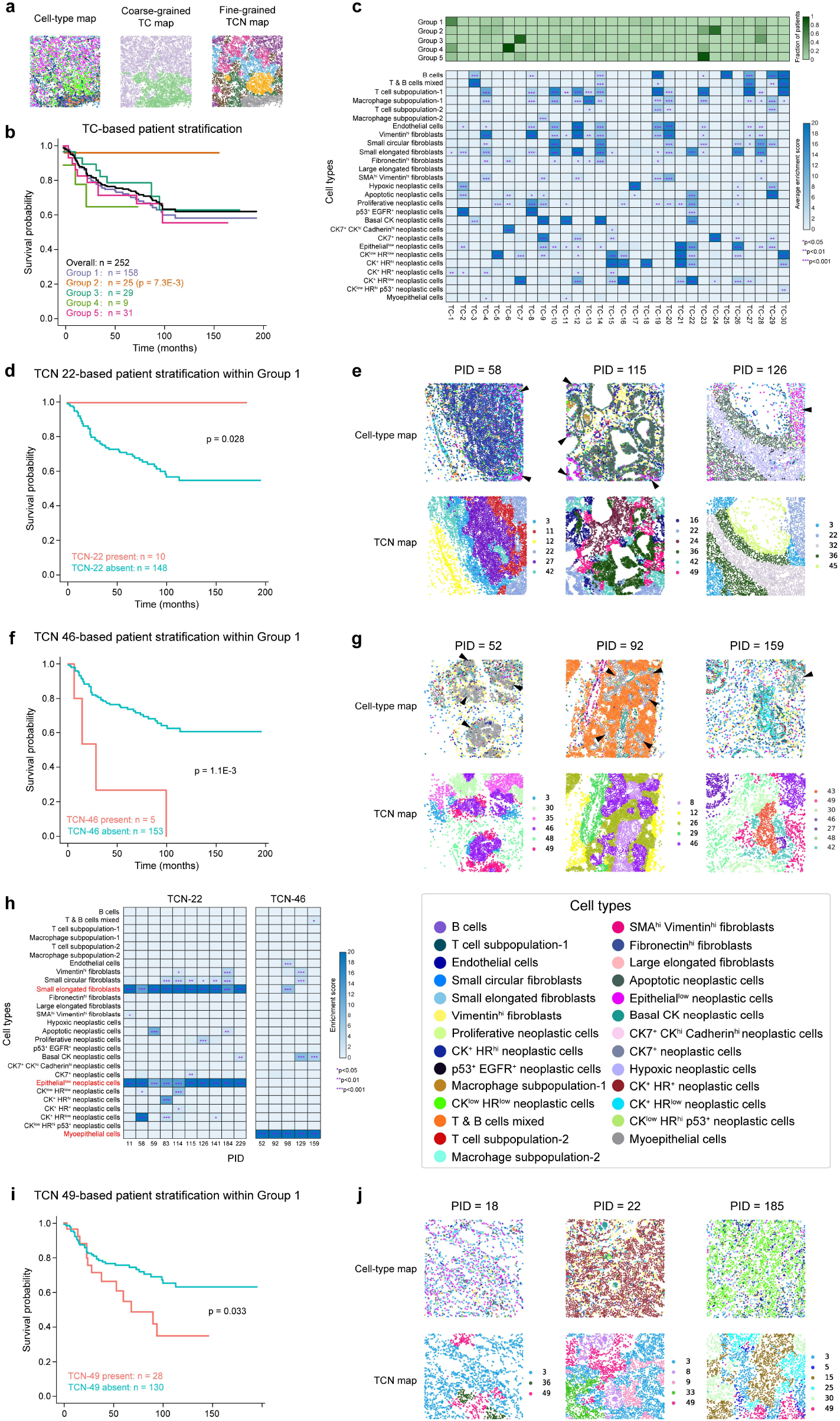
Hierarchical tissue structure-based stratification of breast cancer patients. **(a)** Single-cell spatial maps of breast cancer patient obtained using IMC technology (Taking patients with “PID=11” as an example). The cells are colored according to cell types (first column), the coarse-grained TCs (second column) and the fine-grained TCNs (third column) identified by HRCHY-CytoCommunity. **(b)** Kaplan-Meier survival curves of 252 breast cancer patients, categorized into five groups based on unified TCs. *P*-value was calculated using the log-rank test. **(c)** Heatmaps showing the fraction of patients in each Group with the presence of each unified TC (top) and average enrichment scores for each cell type within each unified TC (bottom). The enrichment score for each cell type is calculated by −log_10_(*P*-value). *P*-values were derived using the hypergeometric test and subsequently adjusted with Benjamini-Hochberg procedure. **(d)** Kaplan-Meier survival curves of patients within Group 1, categorized into two groups based on unified TCN-22. *P*-value was calculated using the log-rank test. **(e)** Representative single-cell maps of patients containing the unified TCN-22 (indicated by arrowheads). The cells are colored based on cell types (first row) and the unified TCNs (second row). **(f)** Kaplan-Meier survival curves of patients within Group 1, categorized into two groups based on unified TCN-46. *P*-value was calculated using the log-rank test. **(g)** Representative single-cell maps of patients containing the unified TCN-46 (indicated by arrowheads). The cells are colored based on cell types (first row) and the unified TCNs (second row). **(h)** Heatmaps showing the average enrichment scores for each cell type within unified TCN-22 and TCN-46 in each patient. The cell types significantly enriched within unified TCN-22 and TCN-46 are marked in red. The enrichment score for each cell type is calculated by −log_10_(*P*-value). *P*-values were derived using the hypergeometric test and subsequently adjusted with Benjamini-Hochberg procedure. **(i)** Kaplan-Meier survival curves of patients within Group 1, categorized into two groups based on unified TCN-49. *P*-value was calculated using the log-rank test. **(j)** Representative single-cell maps of patients containing the unified TCN-49. The cells are colored based on cell types (first row) and the unified TCNs (second row).

For cross-sample comparative analysis, we calculated and concatenated cell type enrichment matrices for TCs and TCNs in all patient samples, respectively (Fig. 1c). Using the K-means clustering algorithm, we identified 30 unified TCs and 50 unified TCNs for this dataset (Methods). By leveraging the presence or absence of these unified TCs as features, we clustered the 252 patients into five distinct groups using the K-means algorithm and performed survival analysis based on the associated clinical data (Fig.5b). Our analysis revealed that the breast cancer patients in Group 2, characterized by the presence of TC-24 (Fig. 5c, top), exhibited significantly better prognosis than the remaining patient cohorts (p-value = 0.0073), whereas patients in other groups showed no significant prognostic differences. Notably, TC-24 represents a coarse-grained tissue structure significantly enriched with CK7^+^ neoplastic cells (Fig. 5c, bottom). This finding aligns with the original study ^6^, which reported that patients with the CK7^+^ neoplastic cell-enriched multicellular structures had better clinical outcomes.

Next, we utilized the presence or absence of a specific unified fine-grained TCN as the feature to further stratify breast cancer patients within the largest Group 1, which is characterized by the enrichment of CK^+^ HR^+^ neoplastic cells (Fig. 5c).

Within Group 1, we observed that 10 patients containing unified TCN-22 exhibited significantly improved prognosis (p-value = 0.028; Fig. 5d). TCN-22 represents a spatial structure enriched with small elongated fibroblasts and epithelial^low^ neoplastic cells (Fig. 5e, top arrowheads; Fig. 5h, left). Normal fibroblasts are known to exert diverse suppressive functions against cancer initiating and metastatic cells through mechanisms such as direct cell-cell contact, paracrine signaling, and maintenance of extracellular matrix integrity. The loss of these suppressive functions is a critical step in tumor progression ^23^. Furthermore, within Group 1, we identified five patients containing unified TCN-46 who had significantly worse prognosis (p-value = 0.0011; Fig. 5f). TCN-46 is a spatial structure enriched with myoepithelial cells (Fig. 5g, top arrowheads; Fig. 5h, right). Growing evidence suggests that myoepithelial cells play a vital role in the organizational development of the mammary gland, and the loss or functional alteration of these cells is a key event in breast cancer progression ^24^.

Additionally, we found that patients containing unified TCN-49 within Group 1 had a significantly worse prognosis (p-value = 0.033; Fig.5i). Interestingly, TCN-49 in these patients did not show enrichment for any cell types (Fig.5j), suggesting that a highly heterogeneous spatial organization of cell types may be associated with unfavorable patient outcomes.

## Discussion

We present HRCHY-CytoCommunity, a GNN-based method for identifying hierarchical tissue structures based on cell type and their spatial location information. The hierarchical tissue structure identification is modeled as a MinCut-based hierarchical community detection problem on a cell-cell spatial proximity graph with node attributes. Unlike the existing method ^18^, HRCHY-CytoCommunity identifies hierarchical tissue structures from a cellular rather than a gene-based perspective, making it suitable for single-cell spatial omics data with a limited number of features, while ensuring that the hierarchical structures cover all cells within the data. By leveraging differentiable graph pooling and graph pruning, HRCHY-CytoCommunity is capable of simultaneously identifying tissue structures of various hierarchical levels at multiple resolutions and exhibiting clearly nested relationship between them. Furthermore, HRCHY-CytoCommunity possesses the ability to discover structures with spatial discontinuous distribution. Additionally, HRCHY-CytoCommunity employs a hierarchical majority voting strategy to ensure the robustness of the result, while maintaining the unambiguously nested relationship between the hierarchical tissue structures. To extend to cross-sample comparative analysis, HRCHY-CytoCommunity utilizes an additional cell-type enrichment-based clustering module to generate a unified set of nested multicellular structures across all tissue samples, thereby addressing the issue of inconsistent hierarchical tissue structure labels between samples.

HRCHY-CytoCommunity successfully identified coarse-grained TCs of varying scales in the mouse spleen CODEX dataset and the TNBC MIBI-TOF dataset, as well as fine-grained TCNs in the mouse hypothalamic preoptic region MERFISH dataset. This suggests that using cell types as cell features can effectively delineate hierarchical functional units within tissues without being influenced by limited gene or protein expression features. Through survival analysis utilizing hierarchical tissue structures as features, we hierarchically stratified patients in the breast cancer IMC dataset, revealing the clinical relevance of hierarchical tissue structures and inspiring further analysis of tumor hierarchical spatial structures and investigation of related immunotherapeutic approaches.

Like most unsupervised clustering algorithms, for each level of the tissue structures, HRCHY-CytoCommunity needs to pre-define the number of multicellular structures, which may impact the identification of hierarchical tissue structures in the absence of corresponding prior knowledge. Additionally, HRCHY-CytoCommunity requires specification of the number of structural levels in each single-cell spatial map, which may impede the discovery of hierarchical tissue structures with different depths in the same single-cell spatial map.

As single-cell spatial omics technologies continue to advance capturing RNA or protein at higher resolutions, there is an increasing need for computational methods capable of identifying hierarchical spatial structures across varying feature dimensions. Using cell types as initial features, HRCHY-CytoCommunity identifies hierarchical tissue structures in an end-to-end manner and lays the foundation for decoding the organization principles by systematically bridging the spatial hierarchy from individual cells through emergent multicellular modules to integrated tissues or organs.

## Methods

### Soft hierarchical tissue structure assignment

We first construct an undirected KNN graph with node attributes for single-cell spatial omics data. In this graph, each node represents a cell, and the node attribute vector using one-hot encoding indicates its cell type information (Fig.1a). To construct the undirected KNN graph, we calculate the Euclidean distance between nodes (cells) based on their spatial coordinates and connect each node to its K nearest neighboring nodes (excluding itself). Each spatial omics dataset is derived from the same tissue type using identical technology. For the undirected KNN graphs, we define the default value of K as the square root of the average cell count per image in the dataset.

For the undirected KNN graph consisting of *n* nodes, we utilize a GNN model ^25^ with the ReLU activation function to produce a node embedding matrix *Z*^(1)^ ∈ ℝ^*n*×*d*^, as described by the following formulation.

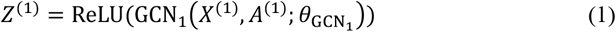

where each row in *Z*^(1)^ is a learned d-dimensional representation vector of each cell. *X*^(1)^ ∈ ℝ^*n*×*m*^ represents the cell type information of all cells, and *m* is the total number of cell types. *A*^(1)^ ∈ {0, 1}^*n*×*n*^ represents the adjacency matrix of the undirected KNN graph. 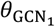 represents trainable parameters in the graph convolutional network. The value of *d* was empirically set to 128 in this study.

Next, we use a fully-connected neural network with no hidden layers (also referred to as a linear layer) and a Softmax activation function to convert the node embedding matrix *Z*^(1)^ ∈ ℝ^*n*×*d*^ into a soft fine-grained tissue structure assignment matrix 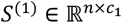, which can be formulated as below.

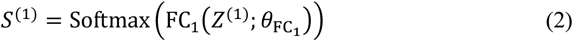

where each element in *S*^(1)^ represents the probability of a cell (row) belonging to one of the *c*_*1*_ fine-grained tissue structures (column). 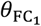 represents trainable parameters in the fully-connected neural network. *c*_*1*_ is a user-specified hyperparameter that indicates the expected number of fine-grained tissue structures to be detected. If the value of *c*_*1*_ is set too large, the deep learning module will automatically determine the optimal number of fine-grained tissue structures, which will be less than *c*_*1*_.

To further identify coarse-grained tissue structures, we utilize a differentiable graph pooling layer ^19, 26^ to generate a coarsened graph with *c*_*1*_ pooled nodes, where each node represents a fine-grained tissue structure. The adjacency matrix 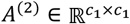and node embedding matrix 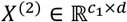 of the coarsened graph are formulated as follows.

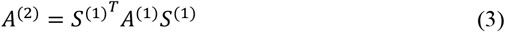

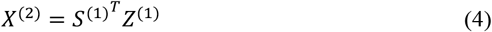

The obtained coarsened graph is fully-connected. The weight of each edge representing the strength of the connection between two fine-grained tissue structures. To relieve the smoothing effect of message passing operation, we introduce a hyperparameter *edge-pruning*, which is empirically set to 0.2. We only retain the edges with weights in the adjacency matrix *A*^(2)^ greater than the *edge-pruning* and reset those edge weights to 1.

Finally, we perform the following graph convolution operation on the coarsened graph to obtain the pooled node updated embedding matrix 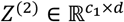, which is then fed into a fully-connected neural network, and the soft coarse-grained tissue structure assignment matrix 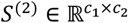 is obtained by a Softmax activation function as below.

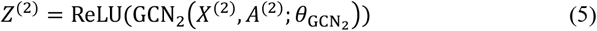

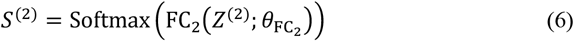

where each element in *S*^(2)^ represents the probability of a fine-grained tissue structure (row) belonging to one of the *c*_*2*_ coarse-grained tissue structures (column). 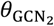 and 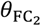 represent trainable parameters in the graph convolutional network and fully-connected neural network, respectively. *c*_*2*_ is a user-specified hyperparameter that indicates the expected number of coarse-grained tissue structures to be detected. If the value of *c*_*2*_ is set too large, the deep learning module will automatically determine the optimal number of coarse-grained tissue structures, which will be less than *c*_*2*_. The overall loss function is defined as follows.

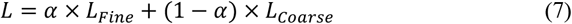

where *α* is a weight parameter used to balance the fine-grained loss *L*_*Fine*_ and the coarse-grained loss *L*_*Coarse*_. Since we prioritize the identification accuracy of fine-grained tissue structures, we empirically set *α* to 0.9. Both *L*_*Fine*_ and *L*_*Coarse*_ are graph MinCut-based loss functions, optimizing the matrix *S*^(1)^ and *S*^(2)^, respectively, which can be formulated as below.

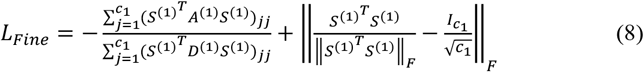

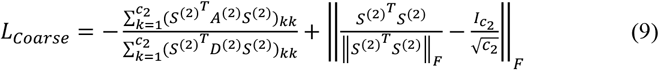

where *D*^(1)^ ∈ ℝ^*n*×*n*^ and 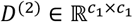 are degree matrices derived from the undirected KNN graph and the coarsened graph, respectively. Both *L*_*Fine*_ and *L*_*Coarse*_ consist of two terms. The left term represents the cut loss term, and minimizing it encourages strongly connected nodes to be clustered together. The right term represents the orthogonality loss term, and minimizing it encourages the tissue structure assignments to be orthogonal and the tissue structures to be of similar size ^19^. In order to prevent the GNN from incorrectly assigning all fine-grained tissue structures to a same coarse-grained tissue structure, we set a threshold *β*, and only retain those results where the orthogonality loss term of *L*_*Coarse*_ is less than *β* after training. We empirically set *β* to 0.7. In order to reduce the scale differences between coarse-grained tissue structures, we set *β* to 0.2 for the TNBC MIBI-TOF dataset, and set *β* to 0.5 for the mouse hypothalamic preoptic region MERFISH dataset. However, if there is prior knowledge indicating a large difference in scale between tissue structures, we should set *β* to 0.7 to avoid the problem of average assignment.

### Hierarchical tissue structure ensemble

After calculating soft hierarchical tissue structure assignment results, we determine the final robust hierarchical tissue structure assignment by the hierarchical tissue structure ensemble module (Fig.1b). First, we run the soft hierarchical tissue structure assignment module several times to obtain multiple soft fine-grained and coarse-grained tissue structure assignment matrices. Subsequently, we multiply the two sets of matrices to obtain a set of matrices representing the probabilities of assigning cells (rows) to coarse-grained tissue structures (columns). Next, for each matrix representing the probabilities that cells assigned to tissue structures, we perform hard assignment by assigning the cell (row) to the tissue structure (column) with the highest probability. Finally, we devise a hierarchical majority-voting strategy described below to integrate the hard fine-grained and coarse-grained tissue structure assignments to obtain a robust hierarchical tissue structure assignment. We empirically set the run times of the soft hierarchical tissue structure assignment module to 20. To reduce the running time, we set the run times to 10 for the spatial proteomics dataset of breast cancer generated using the imaging mass cytometry technology, which contains 252 samples. And we set the run times to five for the mouse spleen CODEX dataset, where each image contains an average of 81,760 cells.

To ensure a clearly nested relationship between coarse-grained and fine-grained tissue structures, we employ a novel hierarchical majority-voting strategy as follows. First, we utilize the majority-voting strategy for all hard fine-grained tissue structure assignments to obtain robust fine-grained tissue structures. Then, we adjust each hard coarse-grained tissue structure assignment by making it incorporate one or several complete fine-grained tissue structures. Finally, we use the majority-voting strategy for all adjusted coarse-grained tissue structure assignments to obtain robust coarse-grained tissue structures.

### Unified hierarchical tissue structure generation across samples

For joint analysis of multiple samples, we align the tissue structures from different samples by a clustering module (Fig.1c). First, we calculate cell type enrichment scores for each tissue structure in each sample using the hypergeometric test. Then, we horizontally concatenate all cell type enrichment score matrices and cluster tissue structures from different samples into a unified tissue structure set. For the TNBC MIBI-TOF dataset, we use the hierarchical clustering algorithm, which facilitates the visualization of the heterogeneity of the 15 patients. For the breast cancer IMC dataset with a large number of samples, we use the K-means clustering algorithm, because it is more efficient.

### Running of NeST

We compared the performance of HRCHY-CytoCommunity to NeST ^18^, the existing state-of-the-art method for identifying nested hierarchical structures in spatial transcriptomics data. In accordance with NeST requirements, we utilized protein or mRNA expression data and cell spatial coordinates as the inputs. For benchmarking purposes, we considered the first layer of the result as the coarse-grained tissue structure and the last layer as the fine-grained tissue structure. In cases where a cell population was assigned to multiple tissue structures within the same layer simultaneously, we allocated them to the tissue structure with the highest accuracy.

The Python code base for running NeST was downloaded from https://github.com/bwalker1/NeST. This code base was applied to three datasets except the spatial proteomics dataset of breast cancer generated using imaging mass cytometry technology. Due to the limited number of features, we adjusted the parameters to obtain several number of single-gene hotspots and hierarchical tissue spatial structures. Specifically, for the mouse spleen CODEX dataset, the parameters were set to be neighbor_eps=400, min_samples=100, hotspot_min_size=100, threshold=0.30, min_size=100, cutoff=0.3, min_genes=2 and resolution=1. For the mouse hypothalamic preoptic region MERFISH dataset, the parameters were set to be neighbor_eps=300, min_samples=2, hotspot_min_size=2, threshold=0.10, min_size=10, cutoff=0.10, min_genes=2 and resolution=1. For the TNBC MIBI-TOF dataset, the parameters were set to be neighbor_eps=100, min_samples=4, hotspot_min_size=5, threshold=0.3, min_size=10, cutoff=0.3, min_genes=2 and resolution=1. Note that NeST does not specify the number of tissue spatial structures identified in advance.

### Quantitative performance evaluation using CODEX and MERFISH datasets

For the mouse spleen CODEX dataset, the ground-truth assignments of cells to fine-grained tissue structures, including red pulp, marginal zone, B-cell zone and periarteriolar lymphoid sheath (PALS), were derived from the original study ^20^. For the mouse hypothalamic preoptic region MERFISH dataset, the outlines of hypothalamic nuclei regions were obtained from the original study ^4^, and the ground-truth assignments of cells to the nuclei were manually annotated based on these outlines. We quantitatively evaluated the performance of HRCHY-CytoCommunity and NeST using three evaluation metrics: Accuracy (ACC), Adjusted Mutual Information (AMI), and Macro-F1 score, with calculations performed using the Python package “scikit-learn (v1.1.2)”. The formulas for these evaluation metrics are as follows.

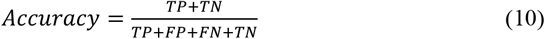

where true positives (TP), true negatives (TN), false positives (FP) and false negatives (FN) represent the comparisons between the predicted tissue structures and the ground-truth cell assignments.

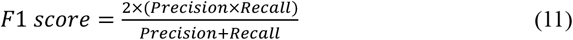

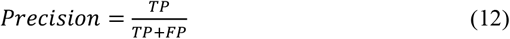

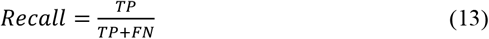

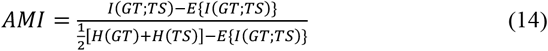

where *E*{*A*(*GT*;*TS*)} denotes the expected mutual information between the ground-truth assignments of cells (GT) and the tissue structure predictions (TS). *H*(*GT*) and *H*(*TS*) represent the entropies of the ground-truth assignments of cells and the tissue structure predictions, respectively. The Macro-F1 score is calculated as the average of the F1 scores across all types of ground-truth tissue structures within the dataset. The AMI is a metric based on Shannon information theory ^27^ that assesses the agreement between the predicted tissue structures and the ground-truth assignments, adjusting for the possibility of random agreement.

### Cell type enrichment score of tissue structures

To perform a quantitative assessment of the cell type composition within tissue structures identified by HRCHY-CytoCommunity, we defined the enrichment score for each cell type in each tissue structure as −log_10_(*P*-value). The *P*-value was determined through a hypergeometric test, taking into account four key parameters: (1) the number of cells of the given type within the tissue structure, (2) the total number of cells within the tissue structure, (3) the number of cells of that type in the image, and (4) the total number of cells in the image. To account for multiple comparisons, the P-values were adjusted using the Benjamini-Hochberg procedure ^28^.

### Survival analysis

In order to verify the ability of HRCHY-CytoCommunity to hierarchically classify patients using hierarchical tissue structures, we took unified TCs and unified TCNs as features and conducted survival analysis combined with the survival data of patients in the breast cancer IMC dataset ^6^. We use the R package “survival (v3.6-4)” to calculate all Kaplan-Meier survival curves and corresponding log-rank test p-values.

## Data availability

This study used the following four publicly available single-cell spatial omics datasets, including a mouse spleen CODEX dataset (https://data.mendeley.com/datasets/zjnpwh8m5b/1), a mouse hypothalamic preoptic region MERFISH dataset (https://datadryad.org/stash/dataset/doi:10.5061/dryad.8t8s248), a human triple-negative breast cancer MIBI-TOF dataset (https://mibi-share.ionpath.com), and a human breast cancer IMC dataset (https://zenodo.org/record/3518284#.Y2UQ0-xBybg).

## Code availability

A software package implementing the HRCHY-CytoCommunity algorithm has been deposited at https://github.com/huBioinfo/HRCHY-CytoCommunity.

## Acknowledgements

This work was supported by a National Natural Science Foundation of China (NSFC) grant No. 62422211 to Yuxuan Hu, NSFC grants No. 62350087, No. 62132015, and No. U22A2037 to Lin Gao, and a NSFC grant No. 62302386 to Jiadong Lin.

We thank Jiadong Lin for helpful discussions. We also thank the Key Laboratory of Computational Bioinformatics of Xi’an at Xidian University for providing computing support.

## Author Contributions Statement

YH and LG conceived and designed the study. ZW, YH and LG designed and ZW implemented the HRCHY-CytoCommunity algorithm. YX provided additional input during method development. ZW performed data analysis. ZW provided support for software package development. YH and LG supervised the overall study. ZW, YH and LG wrote the manuscript.

## Figure Legends

**Supplementary Fig. 1.**
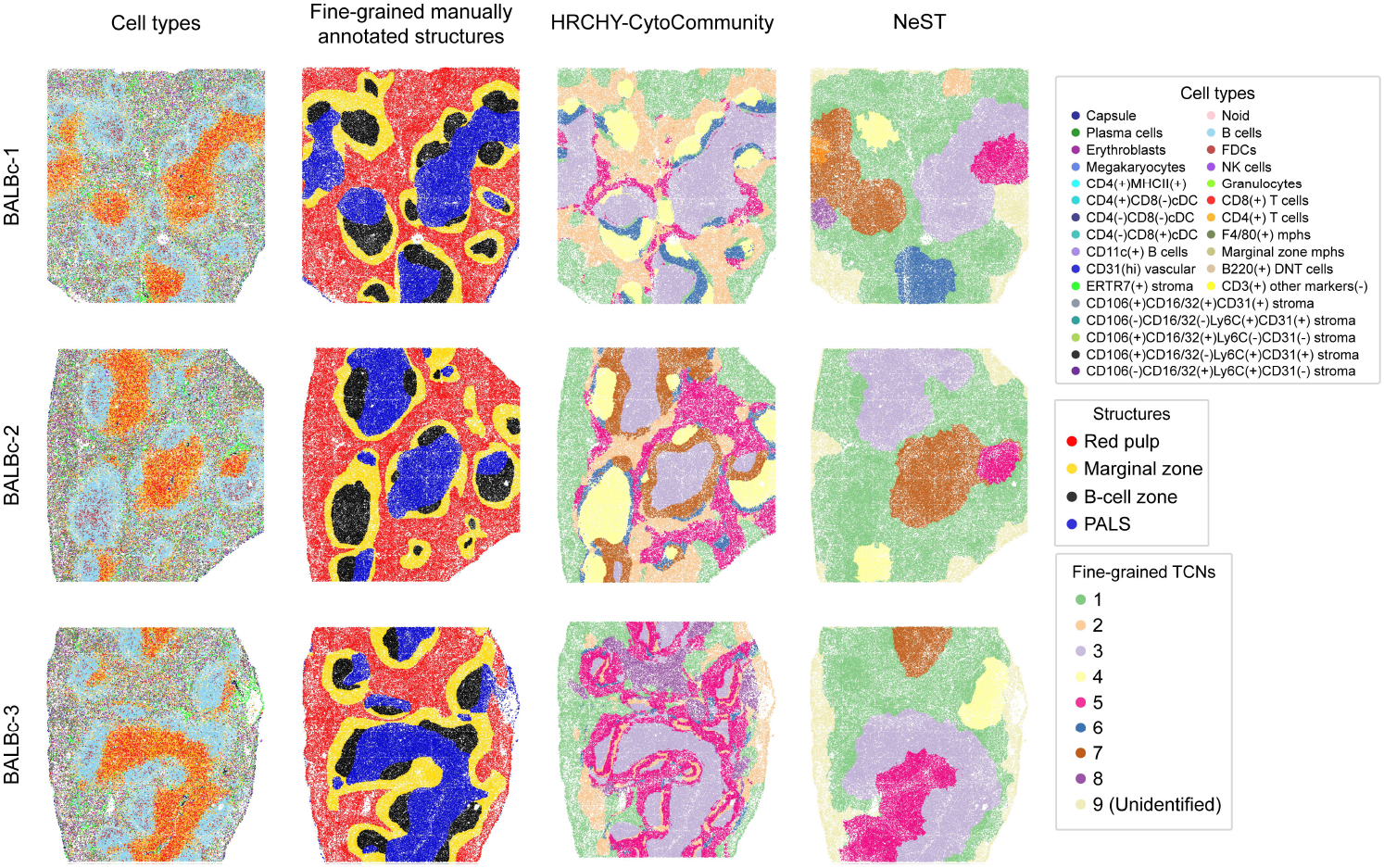
Single-cell spatial maps of healthy mouse spleen. The cells are colored according to cell types (first column), fine-grained manually annotated structures (second column), and the fine-grained TCNs identified by HRCHY-CytoCommunity (third column) and NeST (fourth column). Unidentified indicates that the cells were not assigned to any TCNs.

**Supplementary Fig. 2.**
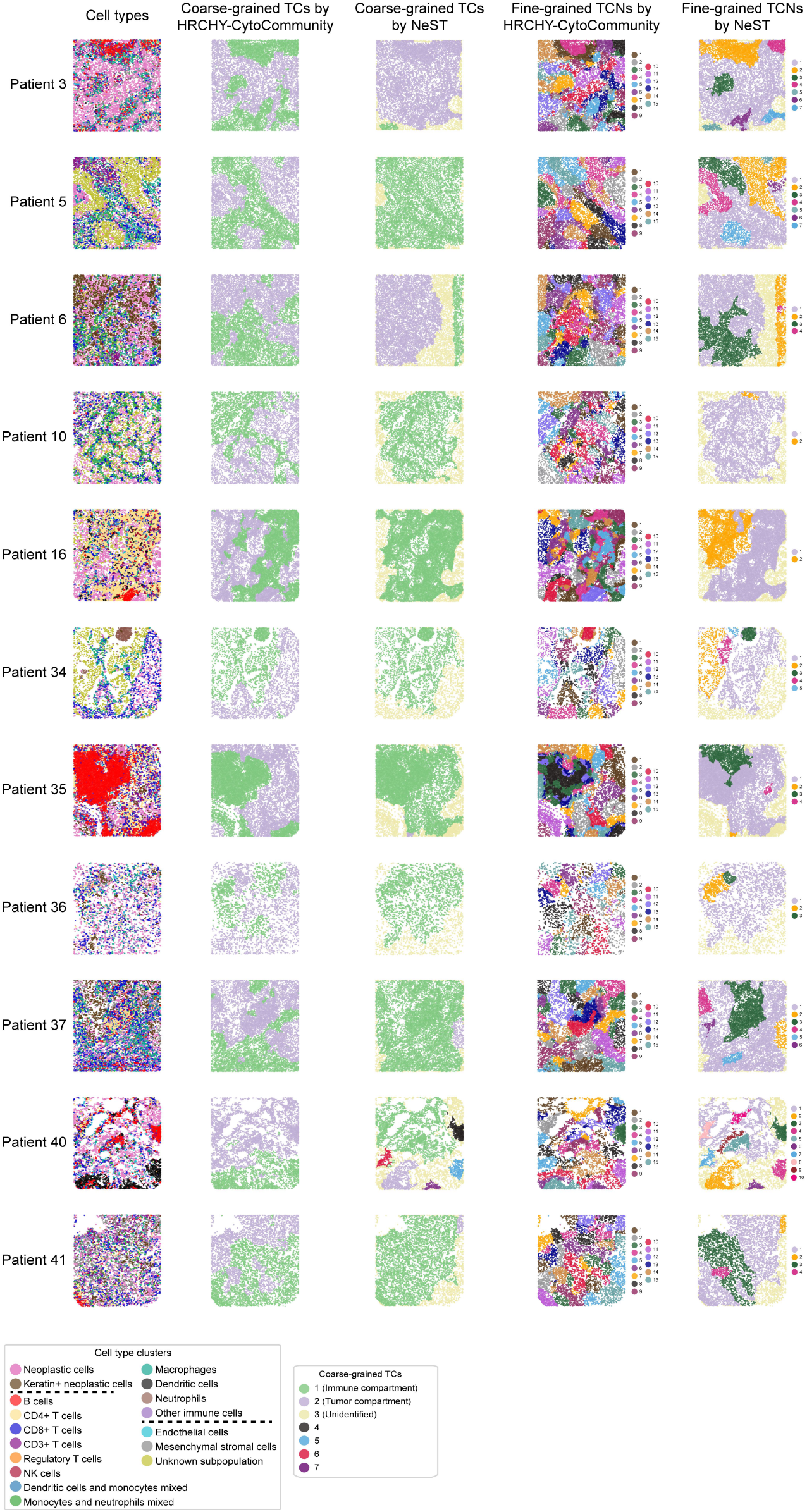
Single-cell spatial maps of the compartmentalized tumors from TNBC patients (except patient 4, 9, 28 and 32). The cells are colored according to cell types (first column), the coarse-grained TCs identified by HRCHY-CytoCommunity (second column) and NeST (third column), and the fine-grained TCNs identified by HRCHY-CytoCommunity (fourth column) and NeST (fifth column). Unidentified indicates that the cells were not assigned to any TCs.

**Supplementary Fig. 3.**
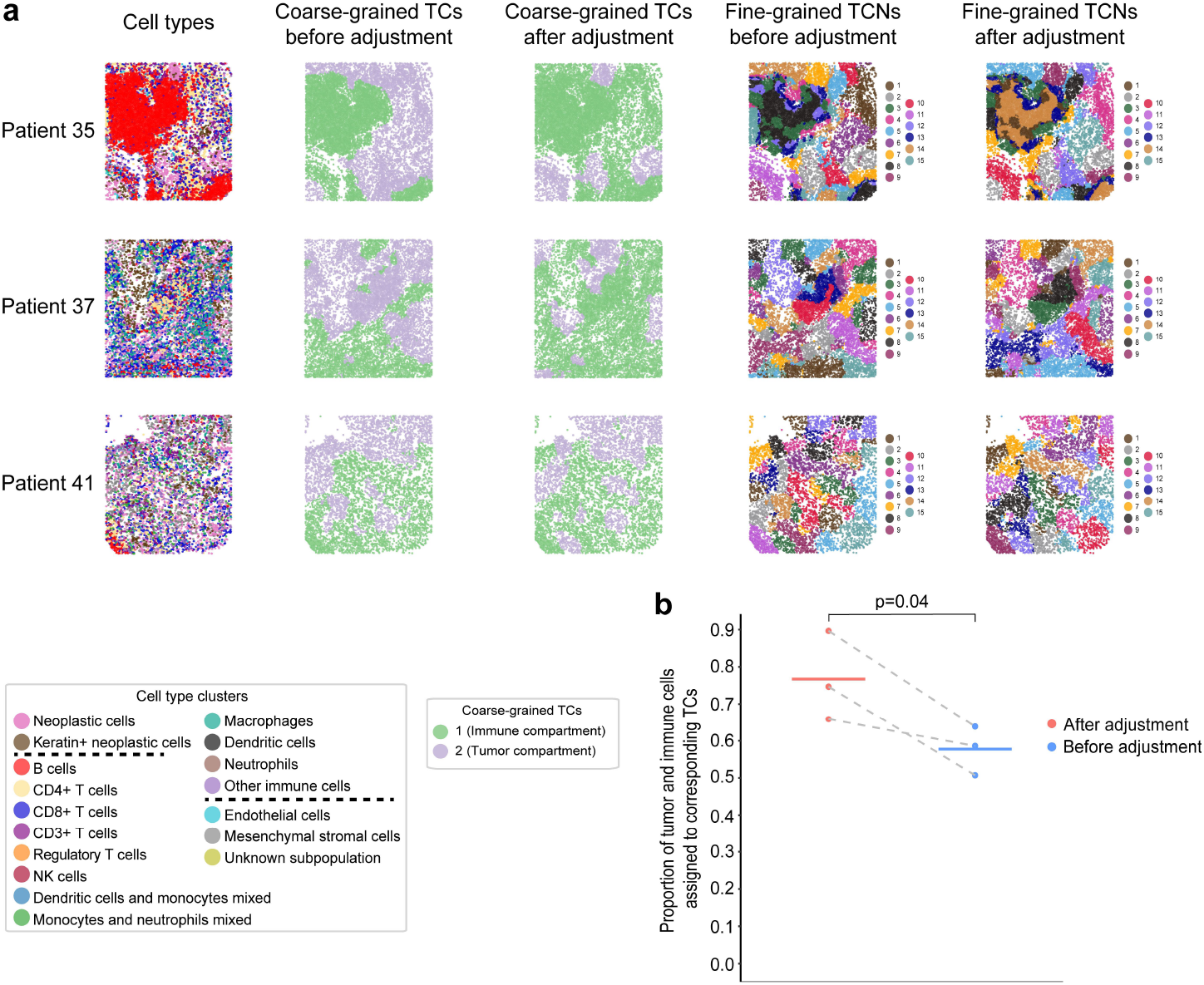
Adjusting the orthogonality loss term threshold of HRCHY-CytoCommunity based on prior knowledge significantly improves the results. **(a)** Single-cell spatial maps of TNBC patient 35, 37 and 41. The cells are colored according to cell types (first column), the coarse-grained TCs identified by HRCHY-CytoCommunity before (second column) and after (third column) adjusting the threshold of the orthogonality loss term, and the fine-grained TCNs identified by HRCHY-CytoCommunity before (fourth column) and after (fifth column) adjusting the threshold of the orthogonality loss term. **(b)** Proportion of neoplastic and immune cells assigned to corresponding TCs. Each point corresponds to the performance on an individual image, with horizontal bars indicating the mean performance across all images. Points from the same images are connected by grey dashed lines. *P*-value was calculated using a one-sided paired t-test.

**Supplementary Fig. 4.**
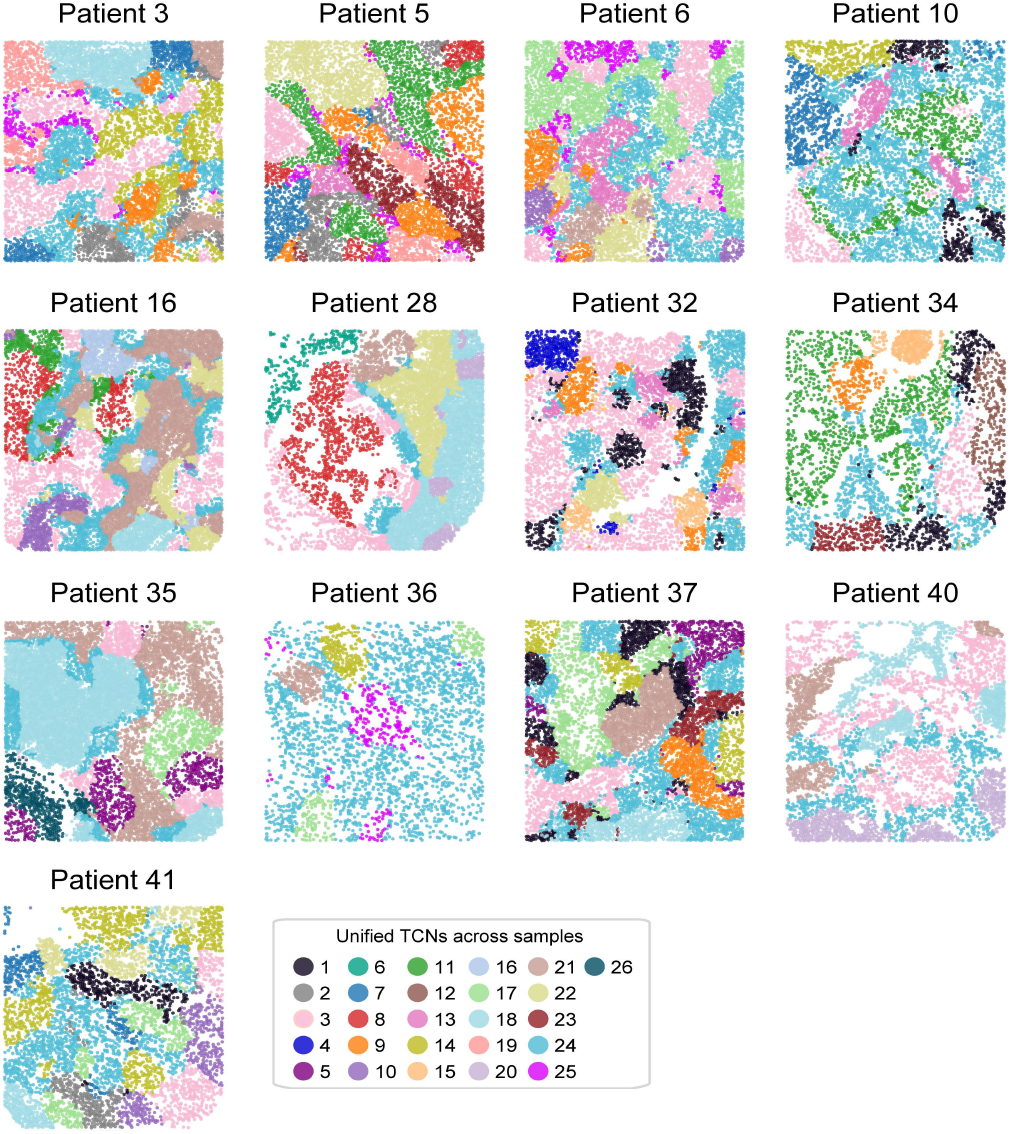
Single-cell spatial maps of TNBC patients except patient 4 and 9. The cells are colored according to the unified TCNs.

## References

1. Ståhl, P.L. et al. Visualization and analysis of gene expression in tissue sections by spatial transcriptomics. Science 353, 78–82 (2016).

2. Rodriques, S.G. et al. Slide-seq: A scalable technology for measuring genome-wide expression at high spatial resolution. Science 363, 1463–1467 (2019).

3. Stickels, R.R. et al. Highly sensitive spatial transcriptomics at near-cellular resolution with Slide-seqV2. Nature Biotechnology 39, 313–319 (2021).

4. Moffitt, J.R. et al. Molecular, spatial, and functional single-cell profiling of the hypothalamic preoptic region. Science 362, eaau5324 (2018).

5. Schürch, C.M. et al. Coordinated cellular neighborhoods orchestrate antitumoral immunity at the colorectal cancer invasive front. Cell 182, 1341–1359.e1319 (2020).

6. Jackson, H.W. et al. The single-cell pathology landscape of breast cancer. Nature 578, 615–620 (2020).

7. Keren, L. et al. A structured tumor-immune microenvironment in triple negative breast cancer revealed by multiplexed ion beam imaging. Cell 174, 1373–1387.e1319 (2018).

8. Rao, A., Barkley, D., França, G.S. & Yanai, I. Exploring tissue architecture using spatial transcriptomics. Nature 596, 211–220 (2021).

9. Zhu, Q., Shah, S., Dries, R., Cai, L. & Yuan, G.-C. Identification of spatially associated subpopulations by combining scRNAseq and sequential fluorescence in situ hybridization data. Nature Biotechnology 36, 1183–1190 (2018).

10. Li, J., Chen, S., Pan, X., Yuan, Y. & Shen, H.-B. Cell clustering for spatial transcriptomics data with graph neural networks. Nature Computational Science 2, 399–408 (2022).

11. Dong, K. & Zhang, S. Deciphering spatial domains from spatially resolved transcriptomics with an adaptive graph attention auto-encoder. Nature Communications 13, 1739 (2022).

12. Long, Y. et al. Spatially informed clustering, integration, and deconvolution of spatial transcriptomics with GraphST. Nature Communications 14, 1155 (2023).

13. Hu, Y. et al. Unsupervised and supervised discovery of tissue cellular neighborhoods from cell phenotypes. Nature Methods 21, 267–278 (2024).

14. Ma, Y. & Zhou, X. Accurate and efficient integrative reference-informed spatial domain detection for spatial transcriptomics. Nature Methods 21, 1231–1244 (2024).

15. Bhate, S.S., Barlow, G.L., Schürch, C.M. & Nolan, G.P. Tissue schematics map the specialization of immune tissue motifs and their appropriation by tumors. Cell Systems 13, 109–130.e106 (2022).

16. Hickey, J.W. et al. Organization of the human intestine at single-cell resolution. Nature 619, 572–584 (2023).

17. Schumacher, T.N. & Thommen, D.S. Tertiary lymphoid structures in cancer. Science 375, eabf9419 (2022).

18. Walker, B.L. & Nie, Q. NeST: nested hierarchical structure identification in spatial transcriptomic data. Nature Communications 14, 6554 (2023).

19. Bianchi, F.M., Grattarola, D. & Alippi, C. in International conference on machine learning 874–883 (PMLR, 2020).

20. Goltsev, Y. et al. Deep profiling of mouse splenic architecture with CODEX multiplexed imaging. Cell 174, 968–981.e915 (2018).

21. Paxinos, G. & Franklin, K.B. Paxinos and Franklin’s the mouse brain in stereotaxic coordinates. (Academic Press, 2019).

22. Chen, R., Peng, B., Zhu, P. & Wang, Y.J.F.i.C.N., Vol. 17 1133445 (Frontiers Media SA, 2023).

23. Alkasalias, T., Moyano-Galceran, L., Arsenian-Henriksson, M. & Lehti, K. Fibroblasts in the tumor microenvironment: shield or spear? International Journal of Molecular Sciences 19, 1532 (2018).

24. Adriance, M.C., Inman, J.L., Petersen, O.W. & Bissell, M.J. Myoepithelial cells: good fences make good neighbors. Breast Cancer Research 7, 1–8 (2005).

25. Morris, C. et al. in Proceedings of the AAAI conference on artificial intelligence, Vol. 33 4602–4609 (2019).

26. Ying, Z. et al. Hierarchical graph representation learning with differentiable pooling. Advances in Neural Information Processing Systems 31 (2018).

27. Romano, S., Vinh, N.X., Bailey, J. & Verspoor, K. Adjusting for chance clustering comparison measures. Journal of Machine Learning Research 17, 1–32 (2016).

28. Benjamini, Y. & Hochberg, Y. Controlling the false discovery rate: a practical and powerful approach to multiple testing. Journal of the Royal Statistical Society: Series B 57, 289–300 (1995).

